# Long-Term Human Skin Platform for Modeling Chronic Inflammation, Environmental Stress, and Therapeutic Intervention

**DOI:** 10.64898/2026.04.30.721933

**Authors:** Purnendu Kumar Sharma, Evelien Schaafsma, Elysia Anderson, Jocelyn Chiari, Paige Thompson, Erin Galt, Sunghoon Lee, Jason Holsapple, Sai Hein, Bailey L Coates, Jade Michaud, Sabrina Zuccaro, Lia Kent, Chris Hinojosa, Kyung-Jin Jang

## Abstract

Chronic inflammation drives tissue dysfunction and aging, yet the dynamic interplay between persistent inflammatory signaling and structural deterioration remains difficult to study in human-relevant systems. Here, an advanced long-term human skin platform is presented that preserves native tissue architecture and epidermal, stromal, and immune-associated molecular programs for up to 4 weeks. Using this system, sustained cytokine-driven inflammation was modeled, demonstrating chronic inflammatory transcriptional programs, progressive histopathological changes, and persistent inflammatory mediator secretion that were broadly suppressed by the JAK inhibitor tofacitinib. Using aged donor tissue, prolonged senolytic-associated treatment attenuated inflammatory and remodeling pathways. Finally, UVB exposure triggered coordinated stress and inflammatory responses that were partially mitigated using topical sunscreen, demonstrating compatibility with environmental stress modeling and topical intervention within preserved tissue architecture. Together, these findings establish a versatile human skin platform for modeling chronic inflammation, aging-associated tissue remodeling, and environmental stress, providing a translational framework for investigating skin tissue dysfunction and evaluating therapeutic interventions.

## 1. Introduction

Chronic, low-grade inflammation is a hallmark of tissue dysfunction and contributes to the progressive decline of organ function across multiple systems [1–3]. This phenomenon, termed inflammaging, arises from cumulative cellular stress and is sustained by mechanisms including cellular senescence, the senescence-associated secretory phenotype (SASP), and age-associated immune dysregulation [4–7]. Although these processes are well characterized at the molecular level, how persistent inflammatory signaling translates into structural tissue deterioration remains incompletely understood. Emerging evidence indicates that resolution of visible inflammation does not necessarily correspond to complete restoration of tissue homeostasis, as residual inflammatory gene expression and tissue-resident immune populations can persist in previously affected sites, highlighting a critical gap in our ability to model these dynamic and relapse-prone states over time [8, 9].

Human skin represents a uniquely accessible and physiologically relevant system to study these processes. As a barrier organ, it integrates intrinsic cellular programs with continuous environmental exposures, functionally linking immune signaling, extracellular matrix (ECM) remodeling, and barrier integrity [10, 11]. During aging, both immune and stromal compartments undergo functional reprogramming toward a pro-inflammatory state, with keratinocyte activation and fibroblast-derived SASP contributing to matrix degradation and barrier dysfunction [12–15]. These processes are further exacerbated by extrinsic stressors such as ultraviolet (UV) radiation, reinforcing the skin as a central model for studying the convergence of chronic inflammation, senescence, and environmental damage [16, 17].

Despite their importance, these complex, multi-layered dynamics remain difficult to study using existing experimental systems. Simplified *in vitro* models lack architectural and cellular complexity, and exhibit altered barrier properties that confound accurate assessment of topical testing [18]. While human skin explants preserve native tissue complexity, they are generally limited to short-term use due to progressive loss of tissue integrity, including the onset of keratinocyte necrosis and epidermal disruption after the first week of culture [19]. Animal models provide valuable insights but they fail to fully recapitulate human-specific immune-stromal interactions and epidermal architecture, limiting their translational relevance [10]. These limitations highlight the need for human-relevant systems that preserve tissue architecture and immune complexity over extended durations, while enabling evaluation of therapeutic strategies targeting complex and chronic tissue conditions [20–22].

Here, we present a long-term human skin platform that preserves native tissue architecture, multicellular interactions, and molecular fidelity for up to 4 weeks in culture. Using this system, we model key dimensions of chronic tissue stress, including sustained cytokine-driven inflammation and its pharmacologic modulation, senolytic-associated treatment responses in aged donor tissue, and environmental stress evaluated via topically applied interventions. This platform provides a robust and translationally relevant framework for dissecting the interplay between inflammatory signaling and structural remodeling in human tissue, and for enabling evaluation of therapeutic strategies targeting chronic skin tissue dysfunction.

## 2. Results

### 2.1. An advanced long-term human skin platform supports extended tissue maintenance

To establish a system suitable for long-term studies of complex human skin biology, we developed a bioengineered *ex vivo* culture platform capable of maintaining native full-thickness human skin tissue over extended durations. The system incorporates a custom 3D-printed insert configured for 24-well plate use, featuring a highly porous gyroid architecture with interconnected pores (∼700 µm pores, ∼70% porosity) that supports skin tissue at an air-liquid interface (ALI) while enabling enhanced nutrient and waste exchange from the basal compartment (Figure 1A and B).

**Figure 1.**
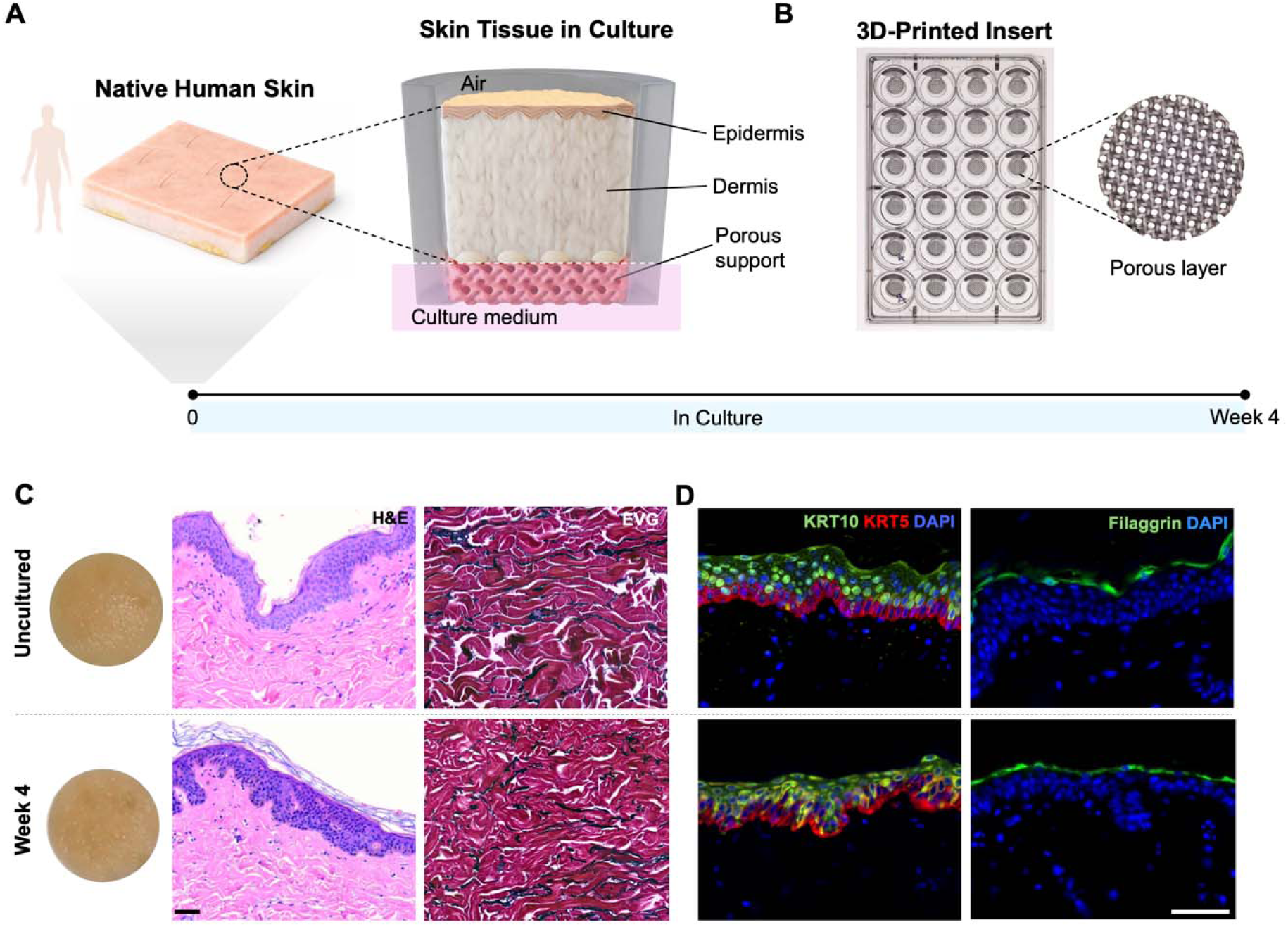
Advanced human skin platform for long-term maintenance of human skin architecture. (A) Schematic of the long-term human skin platform. Full-thickness skin tissues obtained from human donors were processed to generate up to 100 individual biopsies per tissue specimen and maintained at an air-liquid interface on a porous support platform for up to 4 weeks. (B) Top-down photograph of 3D-printed inserts designed to accommodate 8-mm skin tissues in a 24-well plate format. The gyroid porous structure consists of interconnected open pores with an average diameter of ∼700 µm and a porosity of ∼70%. (C) Representative images and histological analysis at uncultured (Day 0) and Week 4 (Day 30). Skin surface photography and H&E and EVG staining were used to assess epidermal and dermal architecture. (D) Representative IF images of epidermal markers over 4 weeks. KRT5 and KRT10 were used to assess basal and suprabasal epidermal organization, and filaggrin was used to assess terminal differentiation and barrier formation. Scale bar, 50 μm.

Human skin tissues were obtained from surgical excision, bulk-processed to generate up to 100 individual biopsies per tissue, and cultured in the platform for up to 4 weeks. No overt macroscopic changes, including discoloration, were observed over time in untreated conditions. Histological assessment demonstrated overall preservation of tissue integrity throughout the culture period (Figure 1C). Hematoxylin and eosin (H&E) and Verhoeff’s Van Gieson (EVG; collagen in red, elastin in black) staining showed that key architectural features were largely retained, including a stratified epidermis, an intact dermal–epidermal interface, and an organized dermal compartment. To further assess epidermal organization, immunofluorescence (IF) staining was performed for markers of basal keratinocytes (keratin 5, KRT5), suprabasal differentiation (keratin 10, KRT10), and terminal differentiation/barrier formation (filaggrin) (Figure 1D). These markers exhibited consistent spatial localization patterns between uncultured (Day 0) and Week 4 samples, indicating preservation of epidermal compartmentalization and differentiation states over time. Together, these findings demonstrate that the platform supports multi-week maintenance of human skin tissue while preserving overall tissue architecture, including epidermal organization and dermal structural integrity, providing a stable foundation for downstream functional and molecular analyses.

### 2.2. Longitudinal transcriptomic analysis reveals stabilization of tissue homeostasis programs

We next characterized the molecular behavior of human skin maintained in the platform over four weeks using bulk RNA sequencing (RNA-seq) across serial time points from 19 donors (Figure 2). Principal component analysis (PCA) showed clear separation between uncultured tissue and cultured samples, consistent with a pronounced early adaptation phase following tissue excision and transport. Notably, samples from Weeks 1–4 occupied a relatively compact transcriptional space rather than diverging progressively over time, indicating an early transcriptional shift followed by stabilization (Figure 2A). To obtain an overview of temporal pathway dynamics, we performed gene set enrichment analysis (GSEA) using Gene Ontology Biological Process (GOBP) terms comparing each culture week to the uncultured condition and highlighted pathways exhibiting distinct temporal patterns, including transient early responses, progressive increases over time, and sustained enrichment across the culture period (Figure 2B). These patterns illustrate dynamic transcriptional adaptations following tissue excision and culture. To more systematically characterize these dynamics, we next identified modules of co-expressed genes with shared temporal trajectories. This analysis shows three dominant patterns corresponding to early adaptive responses, stromal remodeling, and epidermal barrier maturation (Figure 2C). The early adaptive module displayed a sharp transient decline at Week 1 followed by recovery and was enriched for pathways related to structural organization and stress-responsive signaling, including cell adhesion, junction organization, MAPK signaling, and responses to endogenous and osmotic stimuli [23–26]. These changes are consistent with activation of conserved cellular stress response programs triggered by environmental perturbation and macromolecular stress [27]. In contrast, stromal remodeling programs exhibited a progressive increase over time and were enriched for response to growth factor and ECM–associated processes, including ECM regulation, collagen organization, and tissue remodeling. These signatures indicate active restructuring of stromal architecture and dynamic remodeling of matrix components during extended culture. Epidermal differentiation and barrier-associated programs showed early enhancement at Week 1 and remained stable throughout the culture period, including epithelial differentiation, lipid localization, and homeostatic processes, reflecting progressive maturation and stabilization of the epidermal barrier.

**Figure 2.**
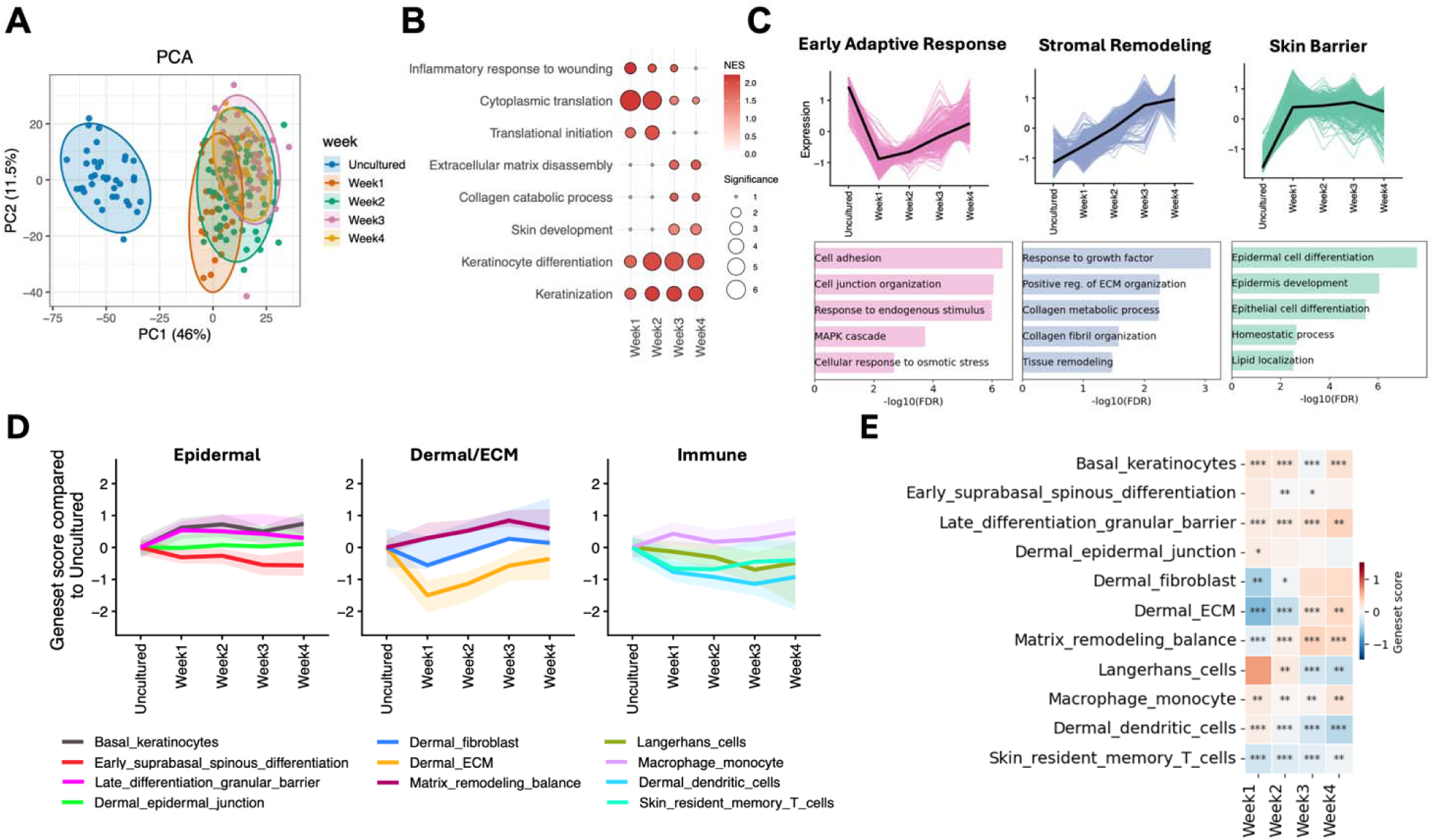
Temporal transcriptomic adaptation of human skin in the long-term platform. (A) PCA of uncultured skin and skin maintained in the platform for 1 to 4 weeks. Data based on 19 donors (118 total replicates). (B) GSEA of GOBP terms comparing each culture week vs. uncultured tissues. Representative significantly enriched upregulated pathways are shown (FDR < 0.05; circle size indicates −log10(FDR), and color indicates NES). (C) Temporal gene expression trajectories of co-expressed genes across time. Colored lines indicate individual genes, and bold black lines indicate median trajectories. Clusters reflect distinct biological programs, including early adaptive response, stromal remodeling, and skin barrier function. Corresponding enriched GOBP terms for each gene cluster are shown below each co-expressed module (FDR < 0.05). (D) Line plot of gene set score differences comparing each culture week to uncultured samples. Gene sets are grouped by compartment: epidermal, dermal/ECM, and immune. (E) Heatmap showing gene set activity scores for the indicated gene sets. Asterisks indicate significant differences between each culture week and the uncultured condition using unpaired t-tests. *P < 0.05, **P < 0.01, ***P < 0.001.

To further contextualize these dynamics at a compartmental level, we next examined compartment- and cell type-associated transcriptional signatures using in-house curated gene sets (Figure 2D and Table S1) [28–35]. In Figure 2D, each signature was anchored to its uncultured baseline to emphasize relative temporal trajectories. Epidermal programs, including basal keratinocyte and late differentiation/barrier-associated signatures, increased over time, whereas early suprabasal differentiation signatures exhibited modest reduction, indicating layer-specific modulation with overall maintenance of epidermal differentiation and maturation. Stromal signatures, including dermal ECM and fibroblast-associated programs, displayed an early decrease followed by recovery over time, accompanied by progressive enrichment of matrix remodeling signatures, consistent with dynamic ECM modulation. Immune-associated signatures showed more modest but cell type–specific temporal shifts, including relatively maintained macrophage/monocyte-associated activity and early detection of Langerhans cell-associated activity. To complement these trajectory plots, Figure 2E displays gene set activity scores in heatmap form, facilitating visualization of gene sets with concordant temporal patterns across culture time points. This representation highlights sustained epidermal differentiation/barrier activity, dynamic matrix remodeling activity, and heterogeneous resident immune-associated transcriptional patterns. Together, these results demonstrate coordinated but non-uniform regulation across epidermal, stromal, and immune compartments over time. These changes reflect ongoing remodeling and a homeostasis-like balance rather than progressive deterioration, consistent with preservation of tissue architecture observed histologically (Figure 1).

### 2.3. Long-term cytokine stimulation induces sustained inflammatory remodeling that is attenuated by tofacitinib

To define the effects of prolonged inflammatory stimulation, we performed longitudinal transcriptomic, histological, and protein-level analyses of IL-17/IL-22-treated human skin tissues over a 3-week period, with or without tofacitinib, a Janus kinase (JAK) inhibitor that suppresses cytokine-dependent inflammatory signaling and has demonstrated efficacy in immune-mediated diseases such as psoriasis [36] (Figure 3A). At Week 1 of treatment, IL-17/IL-22 induced a robust transcriptional response characterized by strong upregulation of inflammatory mediators, including IL36A, DEFB4A/B, S100A12, CXCL1, CXCL8, IL6, and MMP1, alongside suppression of epidermal differentiation and barrier genes such as FLG, FLG2, LORICRIN, KRT77, and DSC1 (Figure 3B). These changes are consistent with cytokine-driven inflammatory activation and impaired barrier function, reflecting established IL-17-driven inflammatory activation and IL-22–associated suppression of keratinocyte differentiation [37–39]. GSEA of the top enriched GOBP terms further revealed enrichment of immune-related pathways, including leukocyte chemotaxis and antimicrobial responses, coupled with suppression of pathways linked to epidermal differentiation and lipid metabolism (Figure 3C). At the pathway level, this profile closely parallels psoriasis-relevant inflammatory transcriptional signatures, characterized by coordinated induction of inflammatory programs and disruption of epidermal differentiation and barrier-associated pathways [37, 39]. Consistent with these findings, hallmark pathway analysis demonstrated sustained activation of inflammatory signaling programs, including TNFα/NF-κB, IL6/JAK/STAT3, and inflammatory response pathways, which were attenuated by tofacitinib [36, 40] (Figure 3D). These findings further support preservation of tissue-resident immune responsiveness over extended culture durations, enabling recapitulation of key inflammatory signaling events in human skin.

**Figure 3.**
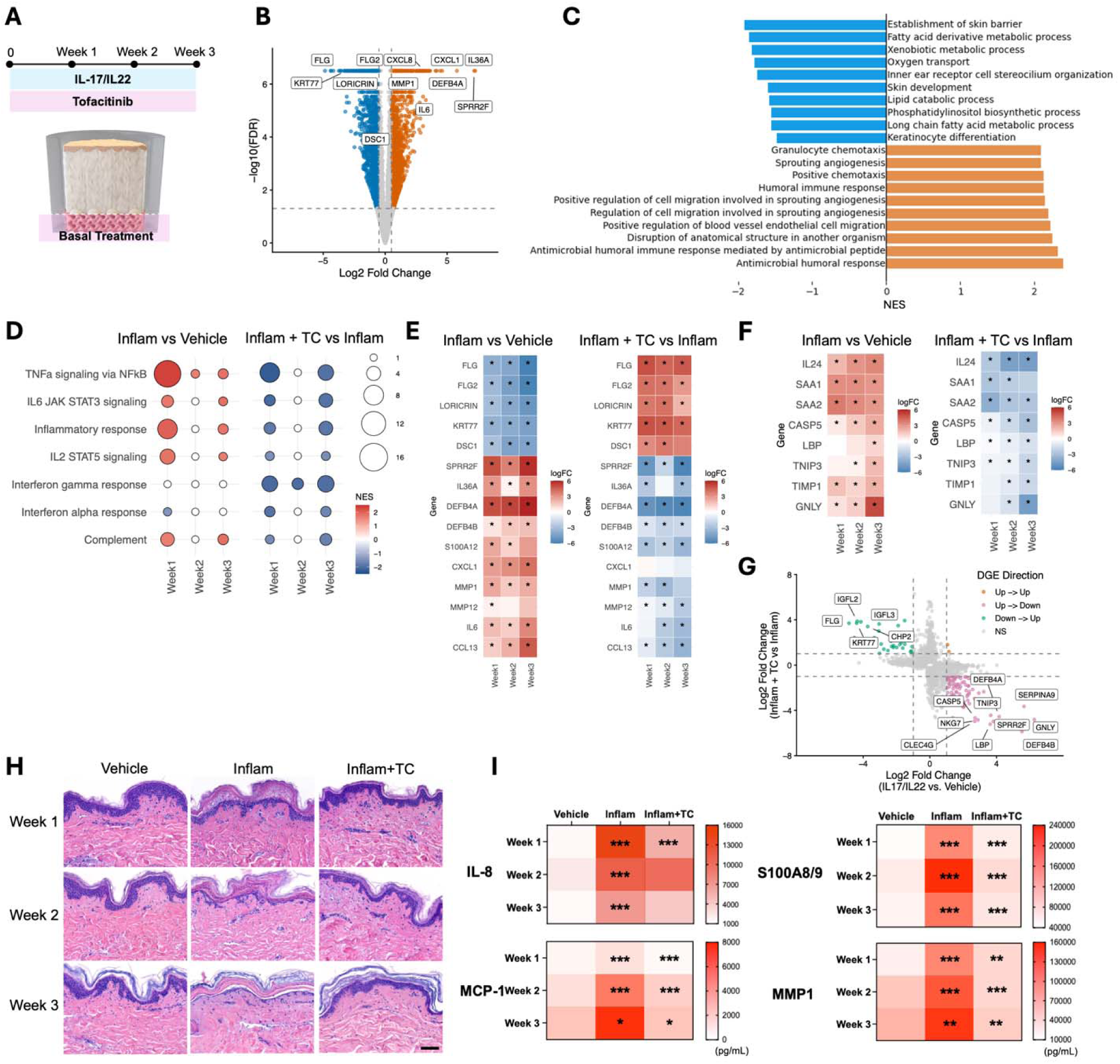
Longitudinal cytokine-driven inflammation and tofacitinib intervention. (A) Schematic of experimental design showing IL-17/IL-22 (Inflam) stimulation with or without tofacitinib (TC) over 3 weeks. (B) Volcano plot of differential gene expression at Week 1 comparing inflamed vs. vehicle-treated tissues. Selected psoriasis-associated marker genes are indicated. Genes passing significant thresholds (FDR < 0.05, |log2 fold change| >0.5) are colored by direction of change: blue, downregulated; orange, upregulated. (C) GSEA of GOBP terms comparing inflamed vs. vehicle-treated tissues. The top significantly enriched upregulated and downregulated pathways are shown (FDR < 0.05). (D) GSEA of Hallmark pathways comparing inflamed vs. vehicle and inflamed + TC vs. inflamed across time points. Circle size indicates −log10(FDR), and color indicates NES. (E) Heatmap of selected differentially expressed genes across vehicle, inflamed, and inflamed + TC over time. Values represent weekly log2 fold changes relative to the vehicle (left heatmap) or inflamed condition (right heatmap). (F) Heatmap of additional chronic inflammation-associated genes across the 3-week time course. Values represent weekly log2 fold changes relative to the vehicle (left heatmap) or inflamed condition (right heatmap). Asterisks indicate a significant difference in gene expression compared to the control group at FDR < 0.05. (G) Transcriptomic reversal plot at Week 3 showing the relationship between gene expression changes induced by inflammation (vs. vehicle) and changes upon TC treatment (inflamed + TC vs. inflamed), based on differentially expressed genes (FDR < 0.05). (H) Representative H&E staining of vehicle, inflamed, and inflamed + TC-treated tissues at Weeks 1, 2, and 3. Scale bar, 100 μm. (I) Heatmap of secreted protein levels (IL-8, MCP-1, S100A8/9, and MMP-1) across treatment groups and time points. Data are presented from 3–4 donors (Week 1 and 2: 4 donors; Week 3: 3 donors; n = 3–9 technical replicates per donor per time point). Multiple comparisons were performed using one-way ANOVA followed by Dunnett’s post hoc test. Asterisks indicate comparisons between inflamed vs. vehicle and inflamed + TC vs. inflamed groups. *P < 0.05, **P < 0.01, ***P < 0.001.

To assess temporal dynamics, we tracked representative inflammatory and barrier-associated genes identified in Figure 3B across the 3-week treatment period (Figure 3E). These genes remained consistently regulated over time, with persistent suppression of barrier-associated genes and sustained induction of inflammatory and remodeling-associated genes across all three weeks. Tofacitinib effectively attenuated both inflammatory gene induction and barrier gene repression across all time points, consistent with inhibition of JAK-dependent signaling components of the cytokine response, particularly IL-22-associated signaling, with secondary suppression of the broader inflammatory program. The sustained responsiveness to tofacitinib further indicates that immune signaling pathways remain active and pharmacologically responsive in the platform over the 3-week period. Prolonged cytokine exposure sustained a set of inflammatory and stress-associated transcripts across the 3-week period, including IL24, SAA1/2, CASP5, LBP, TNIP3, TIMP1, and GNLY [41–44] (Figure 3F). These changes were consistently attenuated by tofacitinib, indicating suppression of both early and sustained components of the inflammatory response. Transcriptomic reversal analysis further demonstrated a shift of IL-17/IL-22-induced genes toward baseline expression following tofacitinib treatment (Figure 3G).

Histological analysis supported these transcriptional findings (Figure 3H). Untreated (data not shown) and vehicle-treated tissues maintained overall epidermal and dermal architecture across all time points. In contrast, IL-17/IL-22-treated tissues exhibited progressive epidermal alterations, including dyskeratosis, basal vacuolar changes, and intracellular edema at Week 1, progressing to parakeratosis and subcorneal separation by Weeks 2 and 3. By Week 3, prominent sloughing of the stratum corneum indicated advanced epidermal barrier disruption. Tofacitinib attenuated these changes, with tissue architecture remaining comparable to vehicle controls and showing minimal epidermal changes. Consistent with these observations, IL-17/IL-22 stimulation induced sustained secretion of IL-8, MCP-1, S100A8/9, and MMP1 across all time points. Tofacitinib broadly attenuated this inflammatory protein response, with significant reductions in MCP-1, S100A8/9, and MMP1 across time points and more modest attenuation of IL-8, consistent with selective inhibition of JAK-dependent cytokine signaling and partial persistence of JAK-independent inflammatory pathways [36, 45] (Figure 3I). Collectively, these data demonstrate that the platform faithfully recapitulates key features of cytokine-driven inflammatory responses observed in human skin, while enabling longitudinal assessment of pharmacologic modulation at transcriptional, structural, and functional levels.

### 2.4. A long-term human skin platform enables evaluation of delayed senolytic-associated responses in aged donor skin

Because senolytic-associated effects often emerge over extended timeframes and are difficult to capture in short-term *in vitro* systems, we next asked whether the platform could elucidate delayed treatment-associated responses in aged tissue. To this end, skin tissues from aged donors (58–74 years) were maintained for three weeks with repeated exposure in the culture medium to vehicle, dasatinib, a tyrosine kinase inhibitor with senolytic activity, or dasatinib plus quercetin (D+Q), a senolytic-associated combination commonly used in aging studies [46, 47] (Figure 4A). Initial analyses at earlier time points showed minimal transcriptional and protein-level responses (Figure S1), consistent with an extended response window over which senolytic-associated effects have been reported to emerge *in vivo* despite the short pharmacokinetic half-lives of these agents [47–49]. Histological assessment at Week 3 showed preservation of overall epidermal and dermal architecture across all treatment groups, with no overt evidence of tissue disruption following prolonged treatment (Figure 4B). At the protein level, both dasatinib and D+Q reduced IL-6 and MMP1 relative to vehicle at Week 3 (Figure 4C).

**Figure 4.**
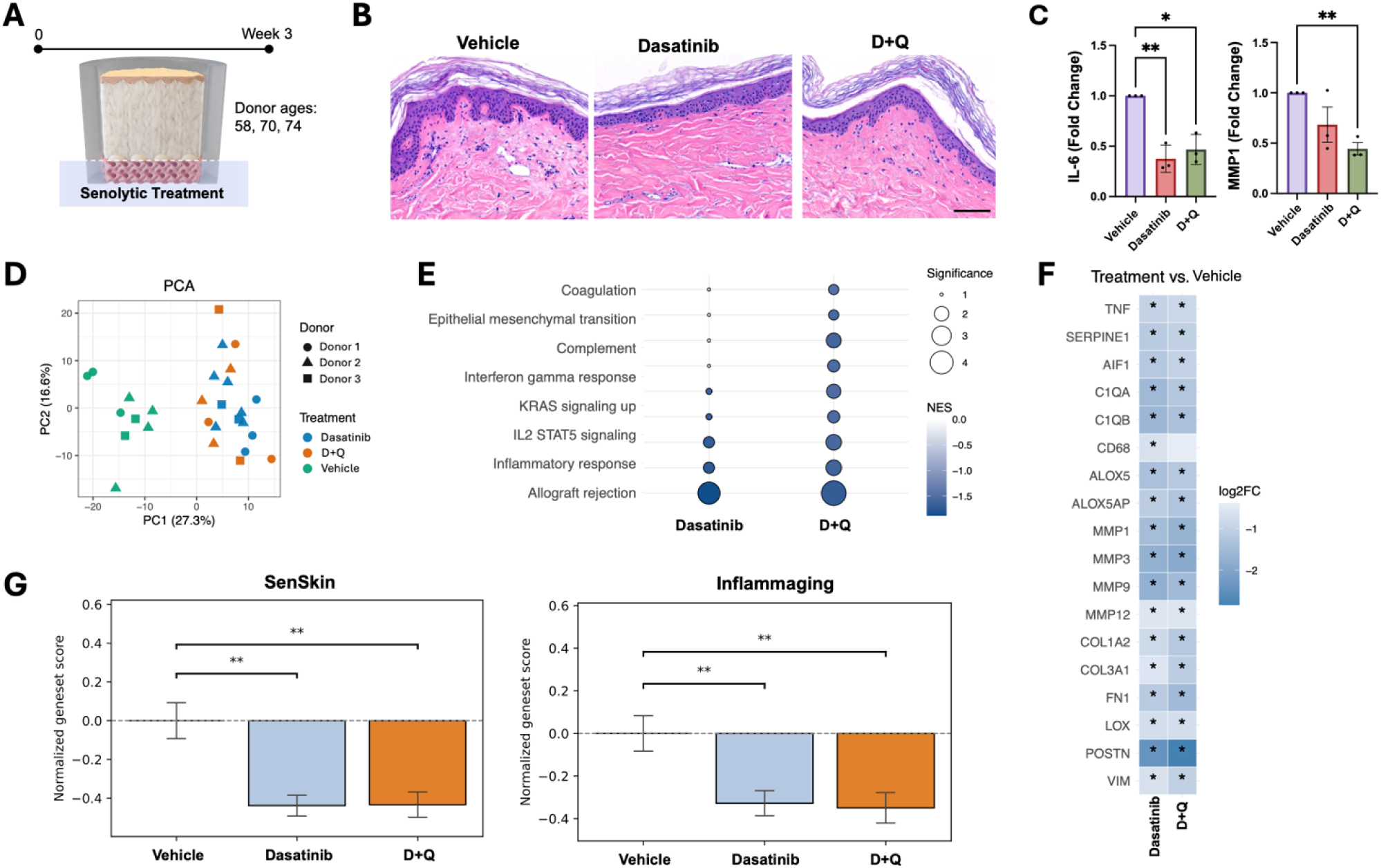
Long-term senolytic treatment in aged skin. (A) Schematic of the experimental design for treatment of aged human skin with dasatinib or dasatinib plus quercetin (D+Q) for 3 weeks. Donor ages are indicated. (B) Representative H&E staining of vehicle-, dasatinib-, and D+Q-treated tissues. Scale bar, 100 μm. (C) Quantification of IL-6 and MMP1 at Week 3 (fold change relative to vehicle control). Data are presented as mean ± SEM from 3 donors (n = 3–5 technical replicates per group per donor). Multiple comparisons vs. vehicle were performed using one-way ANOVA followed by Dunnett’s post hoc test. *P < 0.05, **P < 0.01, ***P < 0.001. (D) Principal component analysis (PCA) plot of transcriptomic profiles stratified by donor and treatment group. (E) Selected pathways from GSEA of Hallmark pathways comparing treatment vs. vehicle. Circle size indicates −log10(FDR), and color indicates NES. (F) Heatmap of differentially expressed genes associated with inflammation and matrix remodeling comparing treatment to vehicle. Values represent log2 fold changes relative to vehicle. Asterisks indicate a significant difference in gene expression compared to the vehicle group at FDR < 0.05. (G) Gene set scores for SenSkin and custom inflammaging signatures across treatment groups relative to vehicle. Statistical significance was assessed using an unpaired t-test for the comparisons shown. Data are represented as median ± SE from 3 donors (n = 2–6 technical replicates per group).

PCA of bulk RNA-seq data revealed separation between vehicle and senolytic-treated samples along the primary principal component, while dasatinib and D+Q largely overlapped, indicating similar transcriptional effects (Figure 4D). GSEA showed coordinated downregulation of immune and inflammatory programs, IL2–STAT5 signaling, interferon gamma response, and inflammatory response, together with suppression of epithelial–mesenchymal transition and related remodeling-associated pathways (Figure 4E). Gene-level analysis supported this pattern, with reduced expression of canonical SASP-associated, inflammatory, and matrix remodeling-related genes (Figure 4F). ALOX5 and ALOX5AP, components of lipid mediator synthesis pathways, were also reduced following treatment [50]. Genes involved in ECM organization and remodeling were modestly downregulated together with matrix-degrading enzymes, consistent with attenuation of active tissue remodeling.

To assess aging-associated transcriptional states, we quantified a published senescence–associated skin gene signature (SenSkin) [51] and an in-house curated inflammaging signature (Table S2). Both dasatinib and D+Q significantly reduced SenSkin and inflammaging scores relative to vehicle (Figure 4G), with broadly similar effects between the two treatments.

Together, these findings demonstrate that the platform enables detection of prolonged senolytic-associated transcriptional responses in human tissue, providing a unique framework to evaluate therapeutic effects that emerge over extended timescales and are difficult to study in short-term *in vitro* systems and conventional experimental models of human aging.

### 2.5. Functional barrier competence for topical intervention: UVB induces epidermal injury responses and inflammatory remodeling

To model UV-induced skin damage and evaluate the platform’s capacity for real-world topical application paradigms–a common limitation of *in vitro* models with altered permeability–human skin tissues were exposed to UVB (300 mJ/cm²) [52, 53] over a 3-day study period in the presence or absence of topically applied sunscreen (Figure 5A). Histological analysis revealed that UVB exposure induced marked epidermal and dermal pathology, including diffuse parakeratosis, dyskeratosis, and hyperkeratosis in the epidermis, as well as focal lymphohistiocytic infiltration and collagen bundle fragmentation in the dermis compared with untreated controls (Figure 5B). These morphological changes were substantially mitigated by sunscreen treatment, which preserved overall tissue structure. Consistent with tissue injury, UVB exposure significantly increased lactate dehydrogenase (LDH) release, indicating cytotoxicity, and elevated secretion of the pro-inflammatory cytokine IL-8 (Figure 5C) in the basal medium.

**Figure 5.**
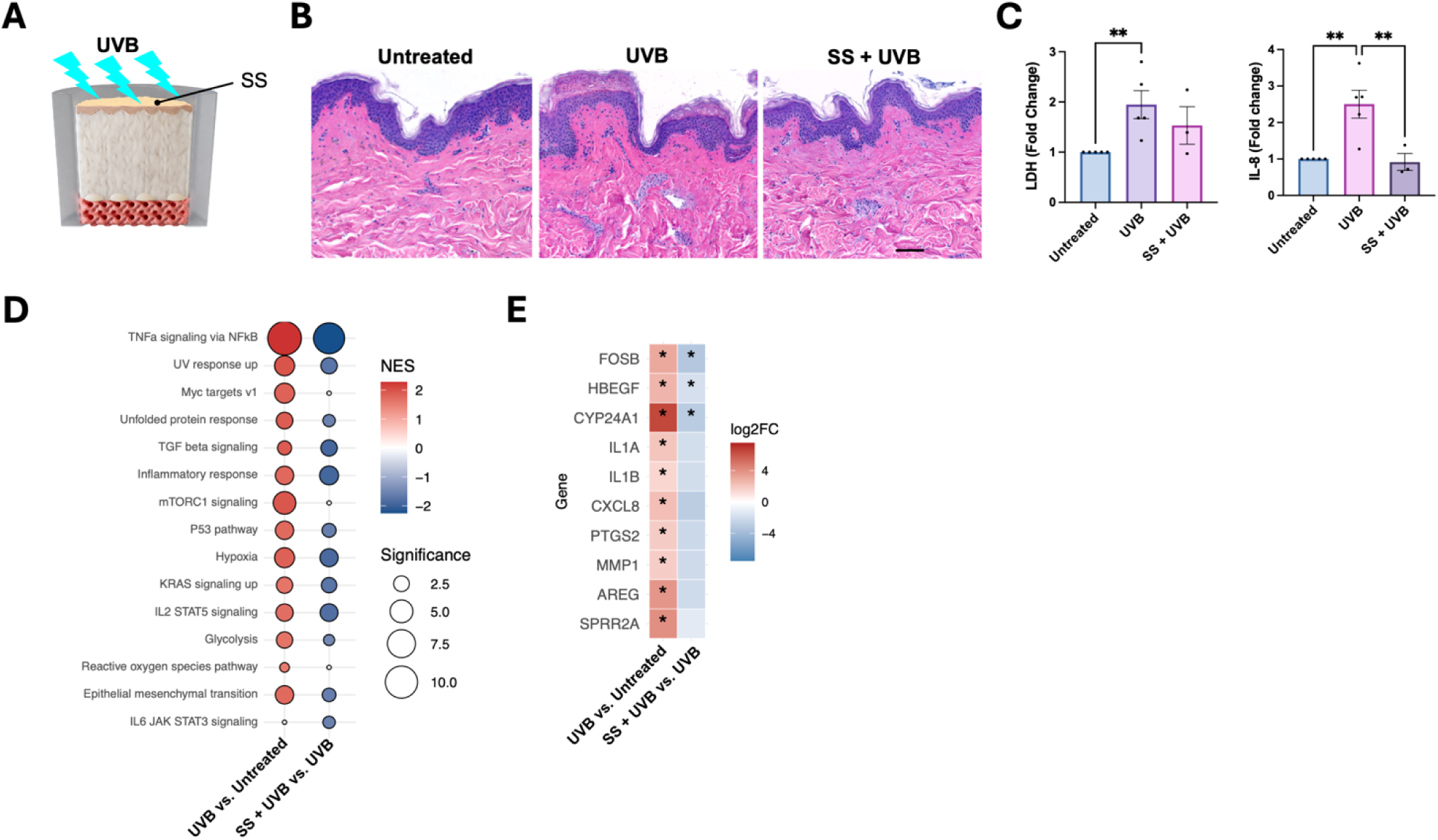
UVB-induced responses and sunscreen protection in the human skin platform. (A) Schematic of the experimental design for UVB exposure (300 mJ/cm^2^) with or without sunscreen (SS). (B) Representative H&E staining of untreated, UVB, and SS + UVB-treated tissues. Scale bar, 100 μm. (C) Quantification of LDH release and IL-8 secretion across treatment groups (fold change relative to untreated control). Data are presented as mean ± SEM from 3–5 donors (n = 3–5 technical replicates per group per donor). Statistical significance was evaluated by one-way ANOVA followed by Dunnett’s post hoc test comparing against the UVB group. Asterisks indicate comparisons between UVB vs. untreated and SS + UVB vs. UVB; *P < 0.05, **P < 0.01, ***P < 0.001. (D) Selected pathways from GSEA of Hallmark pathways comparing UVB vs. untreated and SS + UVB vs. UVB. Circle size indicates −log10(FDR), and color indicates NES. (E) Heatmap of selected differentially expressed genes induced by UVB and suppressed by sunscreen treatment. Values represent log2 fold changes relative to the control indicated on the x-axis labels. Asterisks indicate significant differences in gene expression for the indicated comparison at FDR < 0.05. RNA-seq analyses in panels D and E were performed using samples from one donor.

In the presence of sunscreen, IL-8 levels were significantly reduced and LDH showed a downward trend, suggesting attenuation of UV-induced inflammatory signaling. Sunscreen treatment alone did not induce detectable histological or biochemical changes compared to untreated controls (data not shown).

To elucidate the molecular programs underlying these responses, we performed transcriptomic profiling followed by GSEA. UVB exposure robustly activated pathways associated with inflammatory signaling, cellular stress, and tissue remodeling, including TNFα signaling via NF-κB, UV response, unfolded protein response, and IL6–JAK–STAT3 signaling (Figure 5D). Additional enrichment of MYC, mTORC1, KRAS, and hypoxia pathways indicated broader metabolic and stress-adaptive responses, consistent with *in vivo* UV exposure studies demonstrating temporally structured transcriptional programs, as well as well-established UV-induced inflammatory signaling mediated by ROS, NF-κB, and pro-inflammatory cytokines [54, 55]. These UV-induced pathway signatures were consistently attenuated in sunscreen-treated samples, indicating suppression of stress and inflammatory programs. At the gene level, UVB exposure induced a coordinated transcriptional response encompassing stress-response regulators, inflammatory mediators, and tissue remodeling genes (Figure 5E). Key UV-responsive genes included the AP-1 component FOSB, growth factor signaling molecule HBEGF, and UV-responsive metabolic enzyme CYP24A1, all of which were upregulated following UVB exposure and attenuated by sunscreen treatment. In parallel, canonical inflammatory mediators (IL1A, IL1B, CXCL8, PTGS2) and the matrix degradation-associated gene MMP1 were strongly induced by UVB and showed attenuation with sunscreen. Genes associated with epithelial remodeling and differentiation, such as SPRR2A, also exhibited UV-dependent induction and attenuation with sunscreen (Figure 5E). The ability of the platform to support and evaluate this topical formulation without non-specific toxicity or barrier failure demonstrates its practical compatibility with topical treatment under preserved tissue architecture. Together, these results demonstrate that the platform recapitulates coordinated UV-induced inflammatory and tissue injury responses consistent with *in vivo* human skin, while enabling quantitative evaluation of topical intervention efficacy within a preserved tissue architecture.

## 3. Discussion

The progressive decline of tissue function during aging is widely attributed to “inflammaging”—a chronic, low-grade inflammatory state arising from the interplay of intrinsic and extrinsic stressors [2, 3]. However, resolving how these diverse inputs converge to drive structural tissue deterioration has been limited by the lack of human-relevant models capable of capturing these processes over extended timeframes. Here, we establish a long-term, immune-responsive human skin platform that enables longitudinal interrogation of these dynamics within an intact tissue context. By preserving tissue architecture and sustained immune–stromal signaling for up to 4 weeks, the system maintains key aspects of tissue functionality and provides a tractable framework to investigate chronic tissue stress and aging-associated processes with temporal resolution.

A major technical limitation of *ex vivo* dermatology has been the rapid decline of full-thickness explant viability [19, 56]. Standard transwell systems typically exhibit reduced tissue integrity and loss of functional complexity over time [18, 19]. The platform presented here addresses this limitation through a custom 3D-printed tissue insert with high porosity (∼70%), providing a substantially more open architecture than conventional Transwell membranes and enabling efficient nutrient exchange while maintaining a stable ALI. This configuration supports reproducible, multi-week maintenance of human skin architecture in a scalable SBS-plate format. Longitudinal transcriptomic analysis further revealed that, following an early adaptation phase, tissues stabilize into a differentiated and transcriptionally consistent state rather than undergoing progressive deterioration. Although the system does not recapitulate recruitment of circulating immune cells, it preserves tissue-resident immune and stromal signaling programs over extended culture durations, bridging a key gap between short-term explant models and simplified *in vitro* systems. Histological evidence of lymphocytic infiltration further supports preservation and responsiveness of tissue-resident immune components within the *ex vivo* platform.

A central insight from this model is that distinct biological stressors—including chronic immune activation, senescence-associated signaling, and environmental damage—elicit overlapping features of inflammatory activation, ECM remodeling, and epidermal disruption. These responses were characterized by induction of matrix metalloproteinases, altered keratinocyte differentiation programs, and persistent cytokine signaling, consistent with mechanisms implicated in age-associated tissue dysfunction. Rather than representing independent processes, these findings suggest that diverse stress inputs engage overlapping inflammatory and tissue-remodeling programs, providing a framework for understanding how distinct drivers of tissue stress may contribute to progressive structural decline.

The extended culture duration of the platform enables resolution of distinct temporal kinetics across different stress modalities. Acute environmental insults such as UV exposure induce rapid tissue injury that can be captured over short timeframes, whereas chronic immune activation and aging-associated processes develop over prolonged periods. Cytokine-driven inflammation established a sustained yet pharmacologically reversible state over weeks, while senescence-targeting interventions in aged tissues produced delayed but coordinated attenuation of inflammatory and remodeling-associated transcriptional programs. These delayed responses are consistent with the gradual kinetics reported *in vivo*, highlighting the importance of extended observation windows for evaluating therapies targeting chronic tissue states [47–49].

From a translational perspective, this platform addresses a key gap between conventional *in vitro* systems and *in vivo* human skin biology. Conventional models often fail to capture coordinated interactions between epithelial, stromal, and immune compartments that govern tissue-level responses. By maintaining these interactions within a controlled environment, the system enables integrated assessment of disease-relevant perturbations and therapeutic interventions at molecular, structural, and functional levels. In addition, preservation of stratified epidermal architecture enables evaluation of topically applied interventions within a physiologically relevant barrier context. This is particularly important for dermatologic interventions, for which clinical translation is often constrained by skin penetration, formulation stability, and limited mechanistic validation [57]. The ability to assess UV-induced damage and its modulation by sunscreen therefore demonstrates the platform’s practical compatibility with topical treatment paradigms that remain challenging to model in conventional *in vitro* systems [58, 59].

Several limitations should be considered. While the platform preserves tissue-resident immune components and their associated signaling programs, it does not recapitulate recruitment of circulating immune cells or systemic immune interactions. In addition, the absence of vascular perfusion limits modeling of dynamic nutrient exchange and immune trafficking. Finally, although the use of primary human explants introduces donor-to-donor variability, this feature also enables capture of biologically relevant heterogeneity and may support future studies examining how factors such as age, phototype, and intrinsic skin properties influence nuanced tissue responses.

In summary, this long-term human skin platform provides a versatile system for modeling the interconnected drivers of inflammaging. By enabling investigation of chronic inflammatory signaling, senescence-associated remodeling, and environmental stress within a unified and temporally resolved framework, the model offers a powerful approach for studying mechanisms of tissue aging and evaluating therapeutic strategies targeting chronic tissue dysfunction.

## 4. Conclusion

We establish a long-term *ex vivo* human skin platform that enables multi-week maintenance of native tissue architecture, multicellular composition, and molecular fidelity, supporting longitudinal interrogation of complex tissue dynamics. The system captures key dimensions of chronic tissue stress, including sustained cytokine-driven inflammation, senescence-associated remodeling in aged donor tissue, and UV-induced environmental injury, each eliciting coordinated inflammatory and structural responses that can be modulated pharmacologically or through topical intervention. This platform bridges a critical gap between *in vitro* systems and *in vivo* human skin biology in terms of evaluation duration and human-relevant biological complexity. As such, it provides a versatile and translationally relevant framework for investigating mechanisms of tissue dysfunction and for evaluating therapeutic strategies targeting chronic inflammatory and aging-associated processes.

## 5. Methods

### Insert Manufacturing

Skin biopsy support structures were modeled using Fusion360 (Version 2.0, Autodesk, San Rafael, CA, USA), sliced with PreForm (Version 3.40), and printed on a Formlabs 4B+ printer using Biomed Clear resin (Formlabs Inc., Somerville, MA, USA) at a 50 µm layer height. Parts were washed in 99% isopropyl alcohol for 20 minutes, dried with compressed air, cured for 60 minutes in a Formlabs Cure UV oven at 60 °C, and autoclaved before use.

### Tissue Preparation

Fresh de-identified human skin tissue from healthy donors was procured from both commercial and non-profit suppliers of tissue for research in compliance with collection site IRB protocols and applicable state and federal ethical regulations. Fresh skin tissues, typically measuring approximately 100 cm^2^ in size, were received within 24 hours of surgical removal under controlled conditions. Upon receipt, the tissues were processed under sterile conditions. Full-thickness skin biopsies (8 mm diameter) were generated using custom cutting dies and a pneumatic press (Tippmann, USA). Each sample was trimmed to remove excess subcutaneous fat and underwent a final quality assessment prior to experimental use.

### Tissue Culture

All biopsies were transferred and cultured in sterilized, custom-built 3D-printed insert structures designed to maintain an ALI and enhance nutrient transport. Each biopsy was housed in an individual insert designed to fit in 24 well plates (VWR, Radnor, PA, USA) and supplied with 700 µL of culture media in the basal compartment. The culture medium consisted of a defined formulation based on William’s E medium (Gibco, A12176-01, Thermo Fisher Scientific, Waltham, MA, USA) supplemented with GlutaMAX (Gibco), non-essential amino acids (Gibco), ITS (Corning), Amphotericin B and penicillin-streptomycin (Thermo Fisher Scientific). These cultures were maintained at 37 °C, 5% CO_2_ and 95% humidity. Culture media was replaced every 1–2 days, and effluent was collected for biochemical assays.

### Inflammation Induction and Treatments

Following an initial stabilization period of 48–72 h in culture, tissues from four donors (Donor 1, 36-year-old female; Donor 2, 32-year-old female; Donor 3, 19-year-old female; Donor 4, 52-year-old female) were subjected to inflammatory stimulation using a cytokine cocktail. Biopsies assigned to the inflammation group (Inflam) were treated via the basal media every other day with recombinant human IL-17A (50 ng/mL, BioTechne, 7955-IL-025, Minneapolis, MN, USA) and recombinant human IL-22 (10 ng/mL, BioTechne, 782-IL-010). An anti-inflammatory agent, tofacitinib citrate (TC, 5 µM; MedChemExpress (MCE), HY-40354A, Monmouth Junction, NJ, USA), was administered concurrently with the Inflam group cytokines. Treatments were continued longitudinally for 3 weeks (21 days), and skin biopsies were collected for downstream analysis at each timepoint (week 1, week 2, and week 3). Effluent was collected and replaced with fresh treatment media every other day for a 21-day period (with 3-4 replicates per donor/timepoint). All effluents were stored for biochemical analyses at -80 °C.

### Senolytic treatment

Skin tissue samples from three donors (Donor 1, 74-year-old female; Donor 2, 58-year-old female; Donor 3, 70-year-old female) were stabilized prior to treatment. Tissues were then treated via the basal media with senolytic agents to evaluate their effects on tissue viability and secretory profiles. Treatment groups included Dasatinib (50 nM, MCE, HY-10181) and a combination of Dasatinib and Quercetin (10 µM, MCE, HY-18085). Effluents were collected every other day and replaced with fresh media for all treatments over a 21-day period. All collected effluents were stored for biochemical analyses at -80 °C.

### UV Treatment

Skin tissue samples from five donors (Donor 1, 38-year-old female; Donor 2, 33-year-old female; Donor 3, 21-year-old female; Donor 4, 60-year-old female; Donor 5, 36-year-old female) were exposed to UVB radiation using a UV crosslinker (306 nm; Model 234100, Boekel Scientific, Feasterville, PA, USA). For the irradiation procedure, biopsies from tissue culture plates were transferred to sterile Petri dishes (Greiner Bio-One, 07-000-335, Monroe, NC, USA) containing sterile gauze pads pre-soaked in sterile Dulbecco’s phosphate-buffered saline (D-PBS; Sigma-Aldrich, 56064C, St. Louis, MO, USA) to maintain tissue hydration. These Petri dishes were placed at a fixed distance from the UV source to ensure uniform exposure. Samples were irradiated to a total cumulative dose of 300 mJ/cm² UVB. For the sunscreen treatment group, skin biopsies were pre-treated 15 min prior to UV exposure with a commercially available sunscreen (Neutrogena, SPF 50). A volume of 2 µL (∼4 μL/cm²) was applied to the epidermal surface using a positive displacement pipette (Gilson, Middleton, WI, USA) and was evenly spread using a sterile glass rod. The control samples were handled identically but maintained in culture in the biosafety cabinet for the same duration without UV exposure. Post-exposure, samples were returned to 24-well culture plates containing fresh media and incubated at 37 °C, 5% CO_2_ and 95% humidity. Effluents were collected and replaced with fresh culture media daily.

The study concluded after 3 days, and all effluents were then stored for biochemical analyses at - 80 °C.

### RNA-seq processing and quantification

At each harvest time point, biopsies were bisected perpendicular to the epidermal surface, from the epidermis through the dermis, using a sterile surgical blade (Havalon, 70A, Cincinnati, OH, USA) and stored in RNAlater® (Sigma-Aldrich, R0901) at -80 °C. Samples were then shipped on dry ice to Azenta Life Sciences (South Plainfield, NJ, USA) for RNA extraction, library preparation, and next-generation sequencing.

Total RNA was extracted using Qiagen RNeasy Plus Universal Mini kit following manufacturer’s instructions (Qiagen, Hilden, Germany). Libraries were prepared and sequenced on a NovaSeq (Illumina, San Diego, CA, USA) with 2 × 150 bp paired end reads (∼20M reads/sample). Data were processed using nf-core/rnaseq v3.19.0 [60] of the nf-core collection of workflows [61] with default parameters. The pipeline was executed with Nextflow v25.10.4 [62]. Gene annotations were based on GENCODE v38. Quality control metrics generated by the nf-core/rnaseq pipeline were evaluated for low-quality samples. Principal component analysis (PCA) was used to assess sample clustering and identify potential outliers.

### Differential gene expression (DGE) analysis

We adopted the DGE approach outlined in the nf-core/differentialabundance pipeline (v1.5.0) [63] with minor modifications as outlined below. Count matrices were analyzed for differential expression using DESeq2 (v.1.50.2) [64]. The design formula included the primary contrast of interest along with relevant covariates such as batch, culture week, and interaction terms where applicable. Gene length information was incorporated as an assay (avgTxLength) in the DESeqDataSet object to account for gene length differences. Batch effects were removed using the removeBatchEffect function from the limma package (v3.62.2) [65] on stabilizing transformation (VST)-transformed data and the corrected matrix was used for downstream analyses (referred to as “normalized data” throughout the manuscript).

### Gene set enrichment analysis (GSEA)

Genes were ranked based on the log2 fold changes outputted by DESeq2. GSEA was performed using the fgsea package (v1.32.4) [66]. Gene sets were retrieved from the msigdbr package (v.26.1.0) [67] and included the Hallmark and Gene Ontology Biological Process (GOBP) collections. Pathways were required to have a minimum of 15 and a maximum of 500 genes.

### Culturing time clustering analysis

Temporal gene expression clustering was performed using the Mfuzz package (v2.66.0). Genes with low variance (standard deviation≤0.5, calculated based on normalized data) were excluded. To determine the optimal number of clusters, k-means clustering was applied with default parameters except for *nstart = 25* and with k ranging from 2 to 10 on median-aggregated, normalized gene expression data summarized by week and culture time. The within-cluster sum of squares and the average silhouette width (computed using the cluster package (v2.1.8.2)) were used to determine the optimal number of clusters. Before being inputted into the mfuzz function, normalized gene expression values were standardized by gene to have a mean value of zero and a standard deviation of one (z-scoring). The fuzzifier parameter (m) was estimated from the data. Functional enrichment analysis on the genes in each resulting cluster was performed using gseapy (v1.1.11) [68] and the GOBP geneset collection [69].

### Gene set scoring

Custom gene expression signatures for skin-focused gene sets were quantified using a curated set of canonical marker genes (see Table S1). Marker genes for the SenSkin senescence gene signature were obtained from Wyles et al. [51]. Normalized expression values were z-scored across genes to obtain relative expression levels. For each gene set, expression signatures were computed by averaging z-scored expression values across all genes within the set. For gene set score comparisons between groups, the median gene set score of the experimental group was subtracted from the median gene set score of the reference group to obtain the gene set score difference. Statistical significance between groups was assessed using two-sample t-tests. All RNA-seq and downstream transcriptomic analyses were performed using R v4.5.2 and Python v3.14.3.

### Histology

At each harvest time point, sectioned biopsies were fixed in 10% formalin and shipped the same day to iHisto (Salem, MA, USA) for further processing. The fixed tissues were trimmed, processed and embedded in paraffin, followed by sectioning to 5 µm thickness. Sections were then stained with H&E and EVG for histological evaluation. IF staining was performed using primary antibodies against KRT5 (ab52635, Abcam, Cambridge, UK), KRT10 (ab76318, Abcam), and filaggrin (ab221155, Abcam). Whole-slide images were captured using a PANNORAMIC 1000 digital scanner (3DHISTECH Kft., Budapest, Hungary) by the service provider (iHisto).

### Image Analysis

H&E-stained whole-slide images were analyzed using QuPath (version 0.5.1) for histological assessment. IF images were visualized and analyzed for marker expression using SlideViewer (Version 2.9.0.229983, 3DHISTECH Ltd., Budapest, Hungary).

### Biochemical Assays

The cytotoxicity levels in effluents were assessed by measuring LDH release using the CytoTox 96® Non-Radioactive Cytotoxicity Assay kit (Promega, G1780, Madison, WI, USA), according to the manufacturer’s instructions. Based on study design and requirements, effluents were analyzed for human IL-8 (ELH-IL8-5), human IL-6 (ELH-IL6-5), human MCP1 (ELH-MCP1-5), human MMP1 (ELH-MMP1-5) and human S100A8/9 (ELH-S100A8-9-5) using enzyme-linked immunosorbent assay (ELISA) kits (RayBiotech, Inc., Norcross, GA, USA) in accordance with the manufacturer’s protocols. The absorbance at 450 nm was detected using a Synergy H1 multimode microplate reader (Agilent Technologies, Santa Clara, CA, USA) and analyzed using the built-in Gen5 software (Version3.12).

### Statistical Analysis

All statistical analyses and graphs were produced using GraphPad Prism version 11.0.0 for Mac OS X, GraphPad Software, Boston, Massachusetts USA. Data were analyzed using one-way or two-way analysis of variance (ANOVA), as appropriate, followed by Dunnett’s multiple comparisons test to compare treatment groups against the corresponding control. For all these multi-donor experiments, biological replicates (donors) were treated as independent units (n = number of donors), with technical replicates averaged prior to statistical analysis. Multi-donor bar plot data are presented as mean ± standard error of the mean (SEM). Heat maps represent mean values aggregated across replicates from multi-donor experiments and are visualized using a single gradient color scale. Parameters such as the number of donors, time points, technical replicates, precision (mean ± SEM), statistical tests, and significance are reported in each figure’s legend.

## Acknowledgements

The authors thank Dr. Jungyoon Ohn for critical data interpretation, scientific consultation, curation of the inflammaging gene sets, and manuscript review; Dr. Kevin J. Mills for valuable scientific input and manuscript review; and Dr. Chang Gok Woo for histopathological analysis and interpretation; Jose Fernandez-Alcon for data platform infrastructure and histology data integration; and Dr. Stanley O. King II for establishing tissue sourcing network. We acknowledge the use of tissues procured by the National Disease Research Interchange (NDRI) and the NCI Cooperative Human Tissue Network (CHTN).

## Author Contributions

P.K.S. and E.S. contributed equally to this work. K.-J.J. conceived, designed, and supervised the study, interpreted the data, generated the final figures, and wrote the manuscript. P.K.S. led experimental design and execution, with contributions from E.A., J.C., E.G., P.T., S.H., B.L.C., and J.M., and contributed to data visualization. E.S. performed RNA-seq data analysis, generated associated figures, and contributed to writing the Results section. C.H. designed and manufactured the insert platform, optimized the tissue press workflow, and developed the data platform infrastructure. L.K. and S.Z. contributed to early model and application development.

S.L. and J.H. supported experiments, scientific discussion, and analytical logistics. All authors contributed to the Methods section and reviewed the manuscript.

## Data Availability Statement

The data that support the findings of this study are available from the corresponding author upon reasonable request.

---

**Figure S1.**
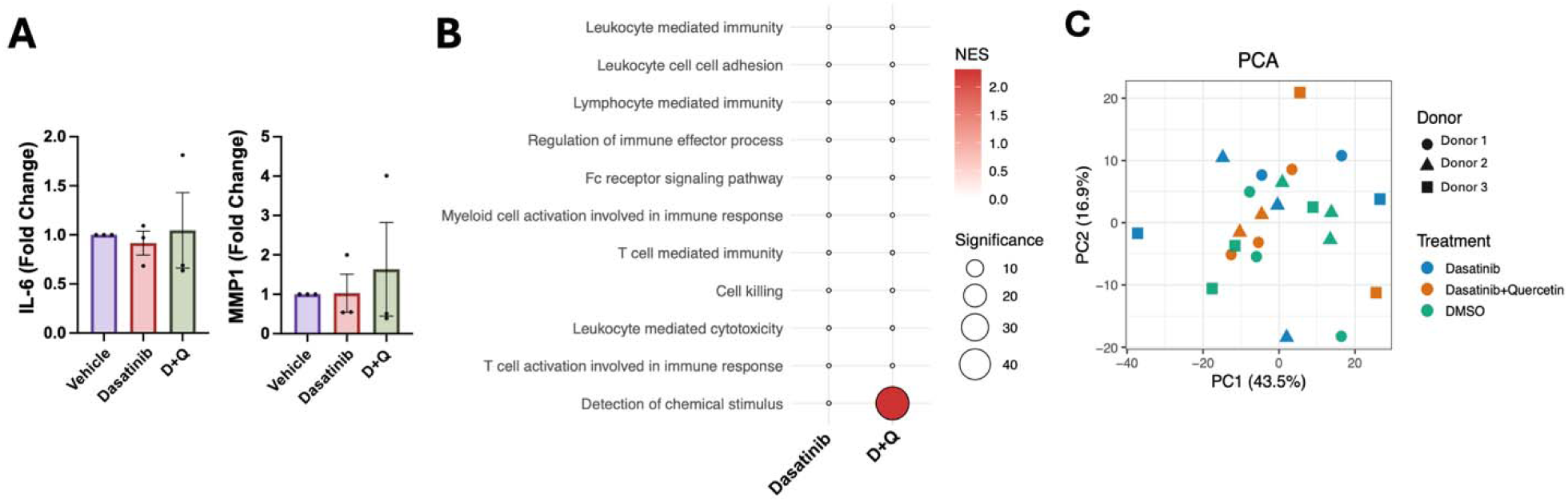
Early senolytic-associated responses in aged human skin. (A) Quantification of IL-6 and MMP1 levels following treatment with vehicle, dasatinib, or dasatinib plus quercetin (D+Q) at Week 1 timepoint. No significant differences were observed between treatment groups. Data are presented as fold change relative to vehicle control. (B) GSEA of selected immune-related pathways comparing treatment groups, showing minimal pathway-level changes at early time points. Circle size indicates −log10(FDR), and color indicates NES. (C) PCA of transcriptomic profiles colored by treatment and donor, indicating no clear separation between groups.

**Table S1.**
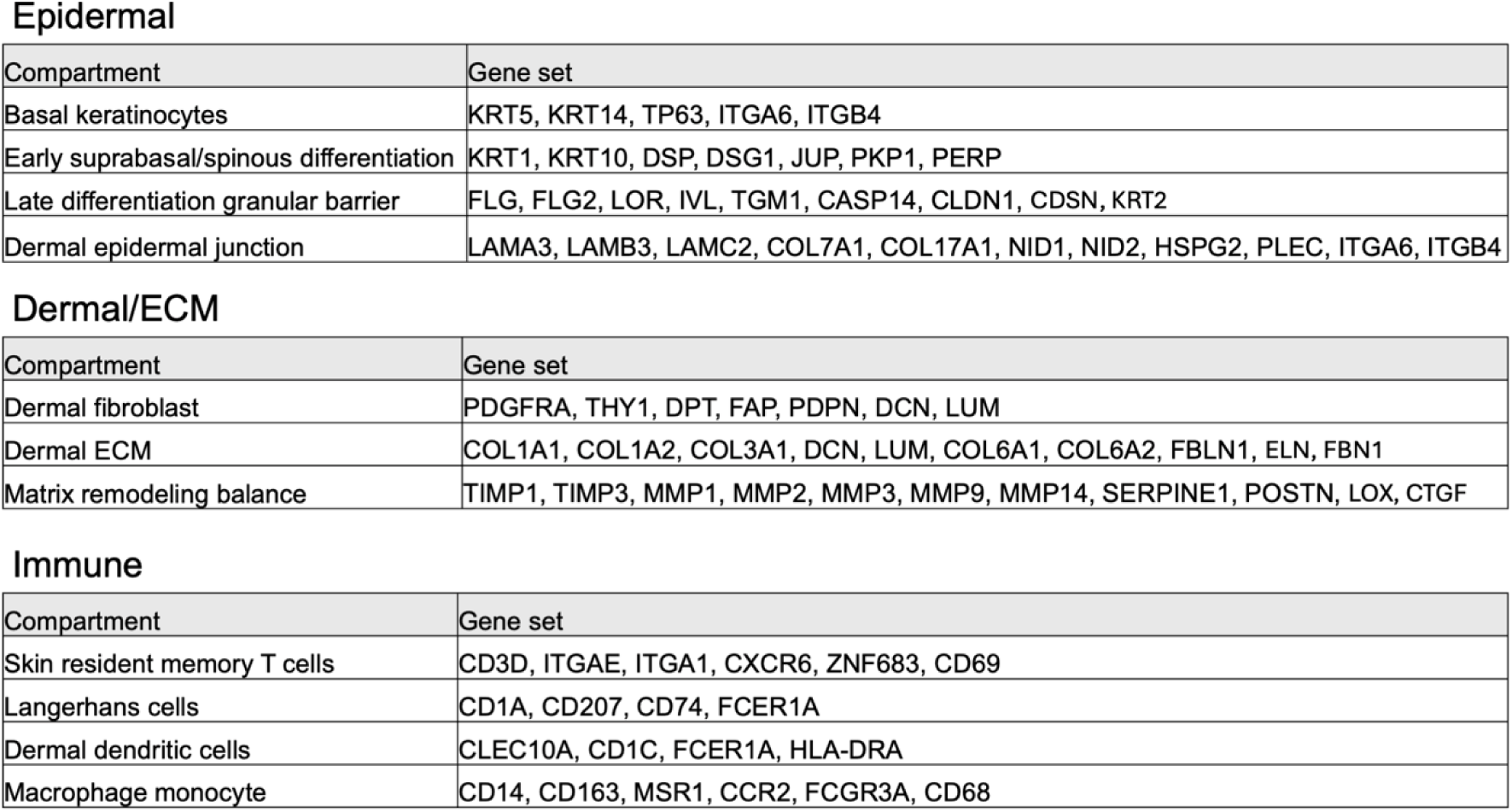
Curated compartment-associated gene sets used for transcriptomic signature analysis. Gene sets representing epidermal, dermal/ECM, and immune-associated compartments used for longitudinal gene set scoring analyses.

Epidermal
| Compartment | Gene set |
| --- | --- |
| Basal keratinocytes | KRT5, KRT14, TP63, ITGA6, ITGB4 |
| Early suprabasal/spinous differentiation | KRT1, KRT10, DSP, DSG1, JUP, PKP1, PERP |
| Late differentiation granular barrier | FLG, FLG2, LOR, IVL, TGM1, CASP14, CLDN1, CDSN, KRT2 |
| Dermal epidermal junction | LAMA3, LAMB3, LAMC2, COL7A1, COL17A1, NID1, NID2, HSPG2, PLEC, ITGA6, ITGB4 |

Dermal/ECM
| Compartment | Gene set |
| --- | --- |
| Dermal fibroblast | PDGFRA, THY1, DPT, FAP, PDPN, DCN, LUM |
| Dermal ECM | COL1A1, COL1A2, COL3A1, DCN, LUM, COL6A1, COL6A2, FBLN1, ELN, FBN1 |
| Matrix remodeling balance | TIMP1, TIMP3, MMP1, MMP2, MMP3, MMP9, MMP14, SERPINE1, POSTN, LOX, CTGF |

**Table S1. Curated compartment-associated gene sets used for transcriptomic signature analysis.**
| Compartment | Gene set |
| --- | --- |
| Skin resident memory T cells | CD3D, ITGAE, ITGA1, CXCR6, ZNF683, CD69 |
| Langerhans cells | CD1A, CD207, CD74, FCER1A |
| Dermal dendritic cells | CLEC10A, CD1C, FCER1A, HLA-DRA |
| Macrophage monocyte | CD14, CD163, MSR1, CCR2, FCGR3A, CD68 |

**Table S2.** Curated inflammaging gene set used for aging-associated transcriptional signature analysis. Inflammaging-associated genes used to quantify chronic inflammatory and aging-associated transcriptional programs in human skin tissues.

|  | Gene set |
| --- | --- |
| Inflammaging | NFKB1, DDX58, IL6, CXCL8, TNF, CCL2, IL18, MMP1, MMP3, MMP9, CTSB, ICAM1, NLRP3, IL1B, IGFBP7, CCN1, CDKN2A, CDKN1A, MTOR, RPS6KB1, NOX4, GDF15, JAK1, STAT3, TMEM173, MITF, TYR, TYRP1, HGF, VEGFA, VCAM1, SERPINE1, C3, CFB, COL1A1, ELN, TIMP1, TGFB1, HAS2, FLG, IVL, LOR, AQP3, DEFB103A, CAMP, CASR, ACLY, SCD, FASN, SIRT1, PPARGC1A, NFE2L2, SOD2, YBX1, LMNB1, JAG1, NOTCH1, IL10, CD163, HMOX1, MMP12, IL1A, AGER, HMGB1, S100A8, S100A9, CXCL12, CDKN1C, IL1A, COL3A1, POSTN, FBN1, MAP1LC3B, PINK1, CLDN1, SPINK5, AREG, CXCL1 |

